# Lack of detectable neoantigen depletion in the untreated cancer genome

**DOI:** 10.1101/478263

**Authors:** Jimmy Van den Eynden, Alejandro Jiménez-Sánchez, Martin L. Miller, Erik Larsson

## Abstract

Somatic mutations in cancer can result in the presentation of mutated peptides on the cell surface, eliciting an immune response. Mutant peptides are presented via HLA molecules and are known as neoantigens. It has been suggested that selection acts against the underlying mutations, leading to neoantigen depletion. Knowing the extent of this specific form of immunoediting may provide fundamental insights into tumour-immune interactions during tumour evolution. Here, we quantified the extent of neoantigen depletion in a wide range of human cancers by studying somatic mutations in the HLA-binding annotated exome, i.e. genomic regions that can be translated into presented peptides. We initially observed reduced non-synonymous mutation rates in presented regions, suggestive of neoantigen depletion. However, when compared to the expected mutation rates from a trinucleotide-based mutational signature model, depletion signals were negligible. This is explained by correlative relationships between the likelihood of mutagenesis in different nucleotide sequences and predicted HLA affinities for corresponding peptides. Our results suggest that signals of immunogenic negative selection are weak or absent in cancer genomics data and that other mechanisms to escape immune responses early during tumour evolution might be more efficient.

## INTRODUCTION

Cancer is caused by somatic mutations in driver genes. These genomic alterations result in a selective growth advantage and positive selection of the affected cells (1). With the rise of next-generation sequencing technologies, increasing insights into the cancer genome have led to a comprehensive characterization of the frequencies and patterns of somatic mutations across different cancers (2, 3). For a tumour to evolve, it also needs to develop ways to avoid immune destruction, a process referred to as immunoediting and one of the more recent hallmarks of cancer (4, 5). Mice studies have shown that T lymphocyte recognition of tumour-specific antigens is crucial for immunoediting to occur (6). The accumulation of somatic mutations in the tumour genome results in the formation of neoantigens, small peptides presented on the cell surface that can stimulate cytotoxic (CD8+) T lymphocytes (CTLs). To attenuate these CTL responses, a cancer cell can upregulate ligands for checkpoint receptors like CTLA4 or PD1. Therapeutically blocking these checkpoint pathways has been shown effective in several cancers like metastatic melanoma and non-small cell lung cancer (7–9). However, responses to immune checkpoint blockade (ICB) therapy are still largely unpredictable and it is not completely understood why some tumours do not respond or develop resistance to therapy.

Several genomic alterations (e.g. *CASP8* mutations, *B2M* mutations, *HLA* loss) have been discovered that can partially explain this ICB therapy unresponsiveness (10–15). Furthermore, as stimulation of CTLs is critically dependent on the formation and presentation of neoantigens, it is not surprising that one of the main determinants of therapy responsiveness is the mutation load (16–18). Indeed, the higher the mutation load, the higher the number of potential neoantigens and hence ways to stimulate the immune system. On the other hand, negative (or purifying) selection is expected to act on mutations leading to neoantigen formation during tumour evolution. This should result in a depletion of neoantigen-forming mutations and escape from immune-induced cancer cell death, potentially even bypassing the need for checkpoint receptor activation. It is hence motivated to determine to what extent neoantigen depletion is detectable in human cancers.

The presence of neoantigen depletion has been suggested in several cancers like colorectal cancer, metastatic melanoma, oesophageal, bladder, cervical and lung cancer (10, 13, 19, 20). As one of the main determinants of CTL immunogenicity is a peptide’s capacity to bind to the cell’s human leukocyte antigens (HLA) from the type I major histocompatibility complex (MHC-I), the conclusions of these studies are based on the demonstration of a lower-than-expected number of non-synonymous somatic mutations in HLA-binding peptides, using the number of synonymous mutations as a reference. Interestingly, it has also been suggested that oncogenic driver mutations, like *BRAF V600E*, preferably occur in patients where the HLA genotype does not lead to the generation of HLA-binding peptides (21).

Somatic mutations are caused by different mutational processes that are active during tumour evolution, resulting in specific mutational signatures in the cancer genome. These signatures are best characterized by the distribution of 96 different trinucleotide substitution types, a combination of the 6 main substitution types and the adjacent up- and downstream nucleotides (3). This implies that the mutation probability at any genomic position is dependent on the immediate sequence context in combination with the active mutational processes, as exemplified by the high frequency of UV-induced TC>TT mutations in metastatic melanoma. It has now been clearly demonstrated that mutational signatures need to be accounted for in any model aiming at finding signals of selection in cancer (22–24). However, it is currently not clear whether and how mutational signatures and their sequence context preferences may have influenced perceived signals of neoantigen depletion in earlier studies.

Here we show that, when these mutational signatures are taken into account, putative signals of neoantigen depletion become very weak to absent in the untreated cancer genome. While our results are in line with the overall weak signals of negative selection in cancer, they challenge the seemingly generally accepted idea about neoantigen depletion shaping the evolving cancer genome, suggesting that other immune evasion mechanisms might be more efficient early during tumour evolution.

## MATERIAL AND METHODS

### Mutation data

MuTect2-called whole exome sequencing (WES) mutation annotation format (maf) files from all 33 available cancer types from The Cancer Genome Atlas (TCGA) were downloaded from the Genomic Data Commons (GDC) Data Portal (data release v7). Colon and rectal adenocarcinoma were considered as a single cancer type for the analysis. All mutation data were fused in a single mutation database and were converted from hg38 to hg19 using UCSC’s liftOver (25). Variants were reannotated using ANNOVAR (26). For each mutation, the main substitution type (i.e. C>A, C>G, C>T, T>A, T>C and T>G) was derived by converting each purine substitution to its complementary base substitution. To determine the trinucleotide substitution type, additional information was added regarding the identity of the upstream and downstream base. Sequence information was derived from UCSC hg19 (25).

### HLA typing

HLA typing of all TCGA samples was performed using Polysolver (11). WES normal bam files from all available TCGA samples were accessed using FireCloud (27), the HLA regions from the main HLA-alleles (HLA-A, HLA-B and HLA-C) in chromosome 6 (coordinates 6:29909037-29913661; 6:31321649-31324964; 6:31236526-31239869) were extracted and the resulting bam files were downloaded. Polysolver was run on these bam files using default settings and without setting prior population probabilities. To validate this HLA typing, the derived frequencies for each HLA allele were compared with the allele frequencies from a healthy US blood donor population, derived from allele frequency net (28) (Fig. S1A).

### HLA affinity predictions and annotation of the HLA-binding genome

Using the R *GenomicRanges* package (29), a *GPos* object was created containing information about the complete exome. For each coding DNA sequence (CDS) position, the amino acid sequences of the 9 possible translated 9-mers (nonapeptides) were determined. HLA affinities of these nonapeptides were predicted for the most frequent HLA alleles (A02:01, A01:01, B07:02, B08:01, C07:01, C07:02) using netMHCPan3.0 (30). For each CDS position, the best binding peptide (peptide with the lowest predicted Kd value) was determined for each of the 6 HLA alleles. Finally, one aggregated Kd value was calculated using the harmonic mean value of the Kd values of the 6 different peptides (one from each allele) and all genomic regions with aggregated Kd values below 500nM were considered as HLA-binding regions.

### Simulation of somatic mutations

All possible point mutations were determined for 17,992 genes by considering for each CDS positions the three possible substitutions (any nucleotide can be substituted in 3 different nucleotides). ANNOVAR was used to annotate the variants and determine the reference and alternative amino acids of each mutation. This information was added to the higher described *GPos* object

To determine the expected somatic mutation rates in the absence of any selection pressure, a simulated mutation database was created, from a similar size as the TCGA mutation database. To match this simulation database for differences in trinucleotide substitution probabilities, we randomly sampled the observed number of mutations from each corresponding substitution type from the *GPos* object.

### Amino acid analysis

To derive the probability of any substitution type to hit a certain amino acid or class of amino acids, we used the *GPos* object containing all possible mutations and determined the amino acid frequency for each substitution type and separately for synonymous and non-synonymous mutations. Amino acids were grouped in 3 classes: hydrophobic (Gly, Ala, Pro, Val, Leu, Iso, Met, Trp and Phe), polar (Ser, Thr, Tyr, Asn, Gln and Cys) and charged (Lys, Arg, His, Gln and Glu).

### Calculation of the HLA-binding mutation ratio (HBMR) and related metrics

To quantify putative signals of immunogenic selection, we defined an HLA-binding mutation ratio (HBMR):

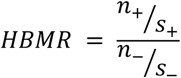

Where n+ and n- are the total number of non-synonymous mutations located in HLA-binding and non-binding regions respectively. Similarly, s+ and s- are the number of synonymous mutations in- and outside HLA-binding genomic regions. A similar metric, called the epitope mutation ratio (EMR) was calculated for the analysis of the IEDB epitopes. Here, + and – refer to the location in- and outside of epitope mapped regions. HBMR *P* values and 95% confidence intervals were calculated using Fisher’s exact test.

dN/dS was calculated considering differences in specific trinucleotide substitution probabilities between cancer types (22):

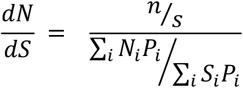

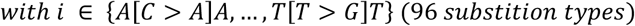

Where N_i_ and S_i_ are the expected number of (non-)synonymous class i substitutions and P_i_ is the probability of substitution class i.

The normalized HBMR was calculated as follows:

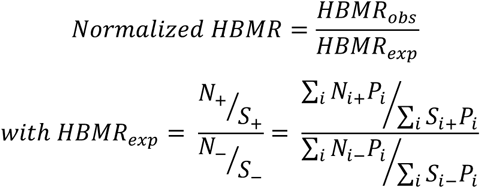

Where N_i+_ and S_i+_ are the expected number of (non-)synonymous class i substitutions in HLA-binding regions, N_i-_ and S_i-_ are the expected number of (non-)synonymous class i substitutions in non-HLA-binding regions respectively and P_i_ is the probability of substitution class i.

### Human epitope mapping

Data from 66,698 known human IEDB (Immune Epitope Database) epitopes were downloaded from synapse (https://www.synapse.org/; id syn11935058) (19). These epitopes were mapped to the human genome (hg19) using the *proteinToGenome* function from the *ensembldb* R package and the *EnsDb.Hsapiens.v75* R library. Mapping was successful for 66,536 (99.8%) epitopes.

### Statistical analysis

The R statistical package was used for all data processing and statistical analysis. Details on statistical tests used are reported in the respective sections.

### Data and software availability

This study is based on public data (open or controlled access) as indicated. The scripts used to produce the results from this manuscript are available on simple request.

## RESULTS

### Annotation of HLA-binding regions in the human genome

Somatic mutations are expected to result in neoantigen formation when 1) the resulting peptides are presented via MHC-I and 2) they are recognized by CTLs through specific T-cell receptor (TCR) binding, which only occurs when there is no immune tolerance, i.e. the presented peptides are new to the immune system. Given sufficient co-stimulatory signals, this neoantigen presentation will result in CTL-mediated cancer cell killing, subjecting the involved mutations to negative selection pressure during tumour evolution (Fig. 1A). We hypothesized that this specific form of negative selection and hence neoantigen depletion should be detectable as reduced mutation rates in genomic regions that can be translated to HLA-binding peptides. Therefore, our first aim was to determine, for each base in the coding exome, whether it is part of a sequence that can be translated into a presented peptide, thereby generating an HLA-binding genomic annotation.

**Figure 1.**
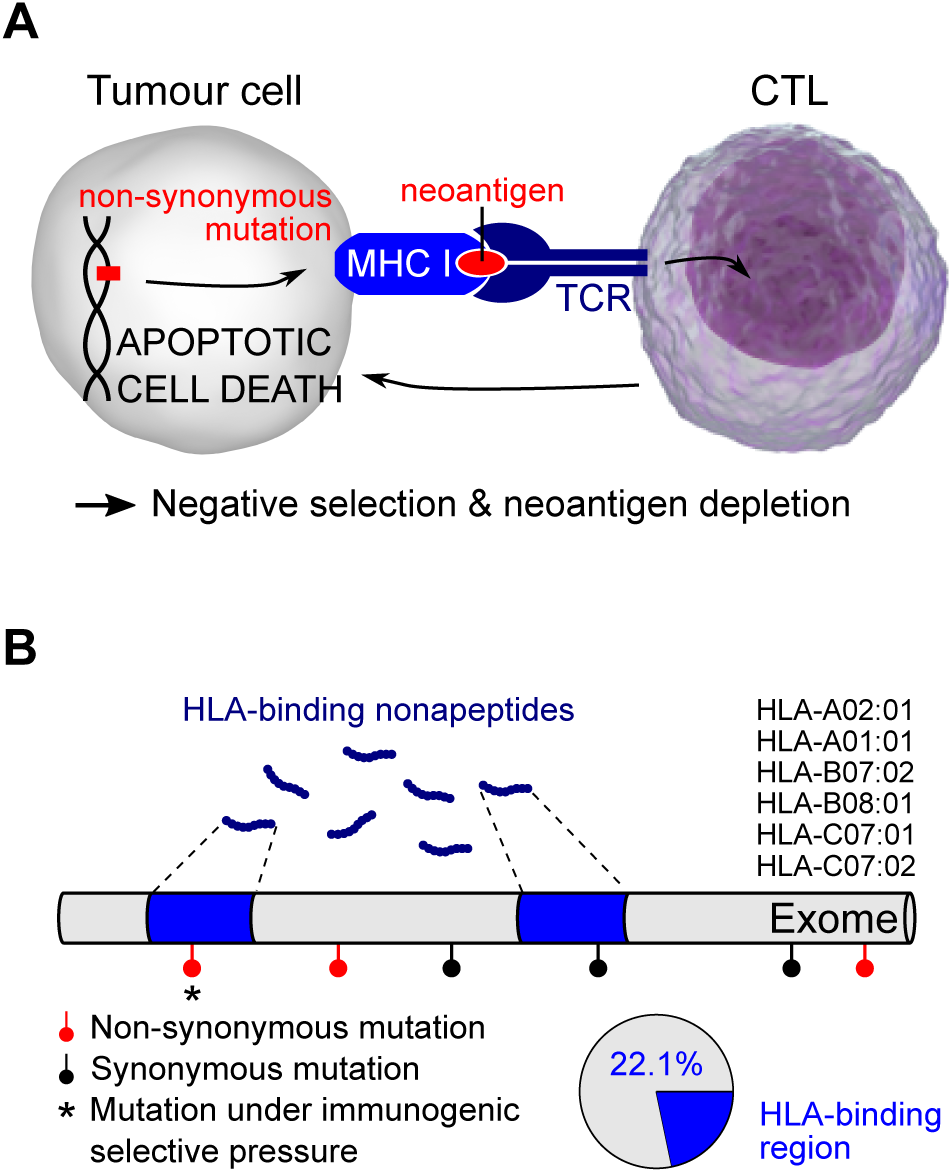
Development of an HLA-binding genomic annotation to detect somatic mutations under immunogenic selective pressure. (**A**) Neoantigen formation is expected when a non-synonymous mutation leads to a structural change in the CTL (CD8+ cytotoxic T lymphocyte) contact residues of an HLA-binding nonapeptide. This will result in CTL-mediated apoptotic cell death and hence negative selection of the underlying somatic mutation. TCR: T cell receptor. MHC-I: type I major histocompatibility complex. (**B**) Binding affinities of all possible nonapeptides were determined for the 6 most common HLA alleles as indicated. Peptides were considered HLA-binding when their aggregated Kd over the 6 alleles was below 500nM (see *Materials and Methods*), HLA-binding peptides mapped to 22.1% of the exome as indicated.

We focused on nonapeptides (9-mers), as these are known to be the most immunogenic (31). HLA-binding affinities of nonapeptides are determined by both the amino acid sequence and by patient-specific HLA genotypes, composed of a combination of 2 HLA-A, 2 HLA-B and 2 HLA-C alleles. We considered a single prototypical HLA genotype consisting of the two most common HLA-alleles (HLA-A01:01, HLA-A02:01 HLA-B07:02, HLA-B08:01, HLA-A07:01 and HLA-C07:02; Fig. S1A), enabling us to define a single HLA-binding genome annotation to use throughout the analyses. The predicted HLA affinities for all nonapeptides translated from the coding genome and for each of these 6 alleles were aggregated in a single affinity, a similar approach to what has been described recently (21) (See *materials and methods* and Fig. S1B-C). By now considering a nonapeptide HLA-binding when the aggregated Kd was lower than 500nM (32), we found that the complete pool of HLA-binding nonapeptides mapped to 22.1% of the exome (Fig. 1B).

### Reduced non-synonymous mutation rates in HLA-binding regions are not caused by selection processes

Having annotated the human exome for the HLA-binding properties of its translated peptides, we next aimed to search for signals of immune-induced negative selection in the cancer genome. All available synonymous and non-synonymous (i.e. missense) somatic mutation data were downloaded from TCGA, encompassing 1,836,369 mutations from 8,683 different samples and spanning 32 cancer types (table S1). Because synonymous mutations cannot change the structure of a presented peptide’s CTL contact residues, only non-synonymous mutations are expected to be under immunogenic selection pressure in HLA-binding regions (Fig. 1A-B). Therefore, we used the number of synonymous mutations as a background reference and estimated the non-synonymous mutation rate based on the ratio of the observed numbers of non-synonymous to synonymous mutations (n/s). Using this ratio, we found lower non-synonymous mutation rates in HLA-binding regions, compared to non-binding regions, on a pan-cancer level (n/s 2.23 in HLA-binding vs. 2.58 in non-binding regions, *P* = 3.24e-198, Fisher’s exact test; Fig. 2A-B). To quantify the extent of this putative neoantigen depletion signal, we defined an HLA-binding mutation ratio (HBMR) as the ratio of the non-synonymous mutation rate (n/s) in HLA-binding to non-binding peptides. This way, immunogenic negative selection of somatic mutations is expected to result in HBMR values lower than 1 (or higher than 1 if these mutations have been influenced by positive selection). For the pan-cancer analysis this implied an HBMR of 0.87, suggesting the overall loss of 13% of non-synonymous mutations due to negative selection (Fig. 2A-B).

**Figure 2.**
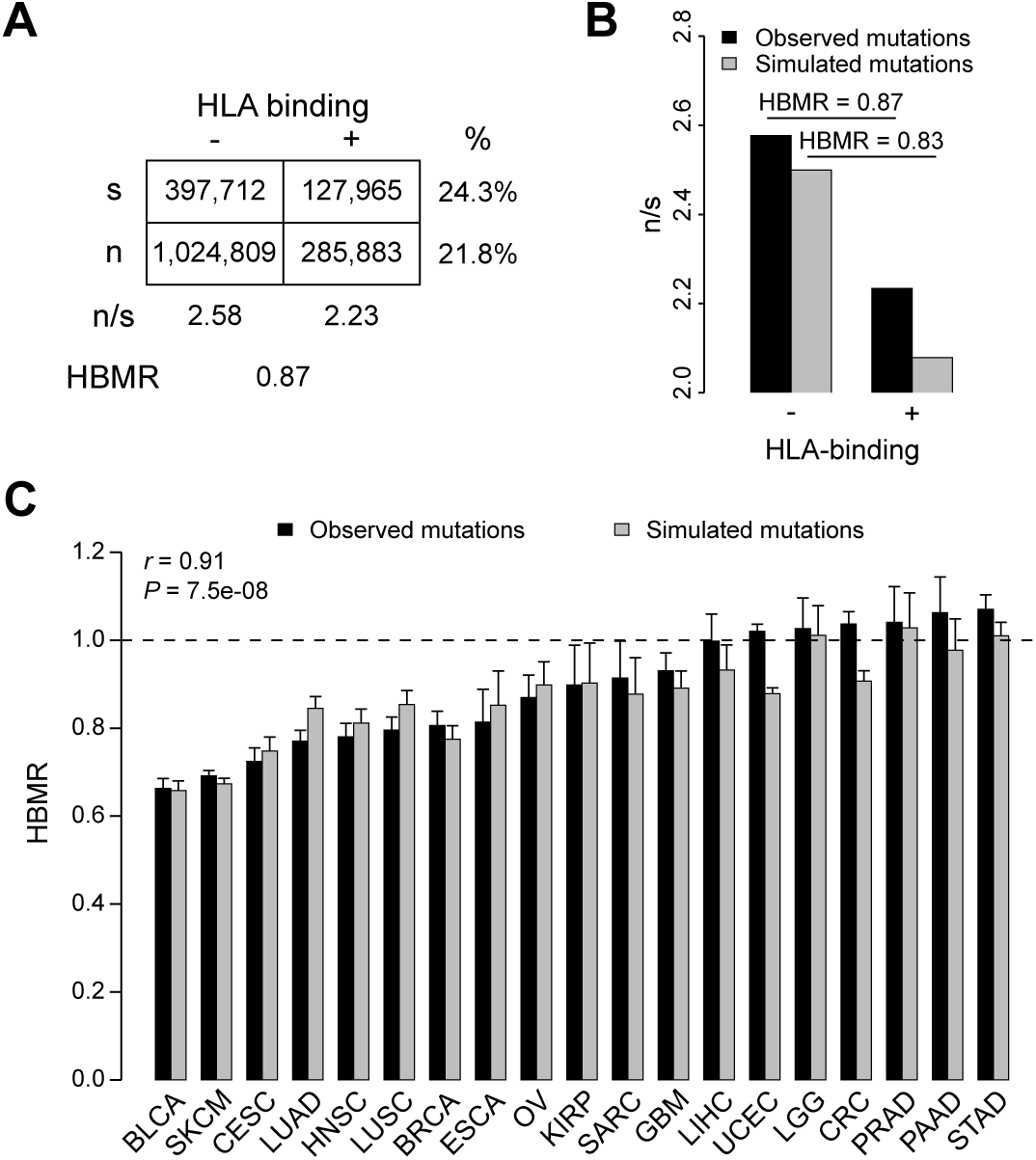
Analysis of somatic mutation rates in HLA-binding annotated genomic regions. (**A**) Table shows the total number of synonymous (s) and non-synonymous (n) mutations, stratified according to their location in the HLA-binding exome. The HLA-binding mutation ratio (HBMR) indicates the ratio of n/s in HLA-binding to non-binding regions. (**B**) Bar plot compares the n/s ratios of observed and simulated mutations. (**C**) HBMR calculated for observed and simulated mutations from 19 cancer types containing at least 10,000 somatic mutations per cancer type. Error bars indicate 95% confidence intervals, calculated using Fisher’s exact test. Pearson correlation coefficient and *P* value indicated on top left. See table S1 for cancer type abbreviations.

We next aimed to determine how these signals differed between cancer types and focused on the 19 cancer types with at least 10,000 mutations in the TCGA dataset. We observed HBMR values that were significantly below 1 for 12 out of 19 analysed cancer types, including bladder cancer (BLCA, HBMR = 0.66, *P* = 1.5e-127), metastatic melanoma (SKCM, HBMR = 0.69, *P* = 0), cervical cancer (…HBMR = 0.72, *P* = 1.3e-51), lung adenocarcinoma (HBMR = 0.77, *P* = 2.3e-60), head and neck cancer (HBMR = 0.78, *P* = 6.6e-36) and squamous cell lung cancer (HBMR = 0.80, *P* = 1.4e-34) (Fig. 2C). In line with these findings, also the dN/dS ratio, which is a widely used metric of selection, was significantly lower than 1 in HLA-binding, but not in non-binding regions, for these cancers (Fig. S2A), again suggesting that negative selection has acted on mutations in these regions as reported earlier (19). We also noted that these findings were robust to the way HLA-binding was determined. Indeed, determining HLA affinities using TCGA sample-specific HLA alleles (rather than the 6 most frequent alleles), pooling affinities of all alleles separately (rather than the aggregated approach) or using a more stringent Kd cut-off of 50nM did not substantially alter the observed reduction in HBMR values (Fig. S3).

To be able to determine whether and to what extent selection processes and hence neoantigen depletion are indeed responsible for the observed reduced mutation rates in HLA-binding regions, we determined the expected mutation rates in the absence of any selection pressure. Therefore, for every observed somatic mutation, we simulated one mutation by randomly sampling from all possible point mutations with the same trinucleotide substitution type (e.g. TCC>TTC), resulting in a simulated mutation dataset from a similar size as the observed data. As expected, all signals of positive selection in driver genes disappeared in the simulated mutation data (Fig. S4). Using this simulated mutation database, we recalculated the mutation rates and HBMR values. Strikingly, a strong signal of apparent negative selection and hence neoantigen depletion, similar to the real mutation data, was still present (HBMR = 0.83, *P* = 0; Fig. 2B). This similarity was also present for the individual cancer types (Pearson correlation coefficient 0.91, *P* = 7.5e-08), with the strongest signals again observed for bladder cancer and metastatic melanoma (Fig. 2C). The fact that a set of randomly generated mutations, upon which selection cannot have acted, gave results that closely mimicked those from actual mutation data casts serious doubt on the interpretation of our results, as well as earlier reports, as signals of neoantigen depletion. As the simulated and real mutations were only matched with respect to trinucleotide substitution types, our interpretation is that different sequence contexts and substitution type probabilities between cancers somehow correlate to HLA-binding properties to create false signals of neoantigen depletion.

### Different trinucleotide substitution probabilities explain lower non-synonymous mutation rates in HLA-binding regions

To better understand how trinucleotide substitution types differ with respect to the probability of affecting a sequence that can be translated to HLA-binding peptides, we determined all possible point mutations in 17,992 genes (21,203,704 synonymous and 67,766,542 non-synonymous mutations; Fig. 3A) and determined how the expected non-synonymous mutation rates differed between HLA-binding and non-binding regions for each trinucleotide substitution type. Using the HBMR to quantify this difference, we noted a remarkable variability between the trinucleotide substitution types, with HBMR values ranging from 0.35 for TCT>TGT substitutions to 2.07 for ATG>ACG substitutions (Fig. 3B). The trinucleotide substitution types with the lowest HBMR values were the most abundant in the cancer types with low overall HBMRs (e.g. 23.9% of all malignant melanoma mutations are TCC>TTC, the trinucleotide substitution type with the second to lowest HBMR; Fig. S5A). Remarkably, many of the substitution types with the lowest HBMR values were TCN>TNN and indeed, a strong negative correlation was observed between the proportion of TCN>TNN mutations in cancer types and their corresponding HBMR values (Pearson correlation r = −0.81, *P* = 2.5e-05; Fig. S5B).

**Figure 3.**
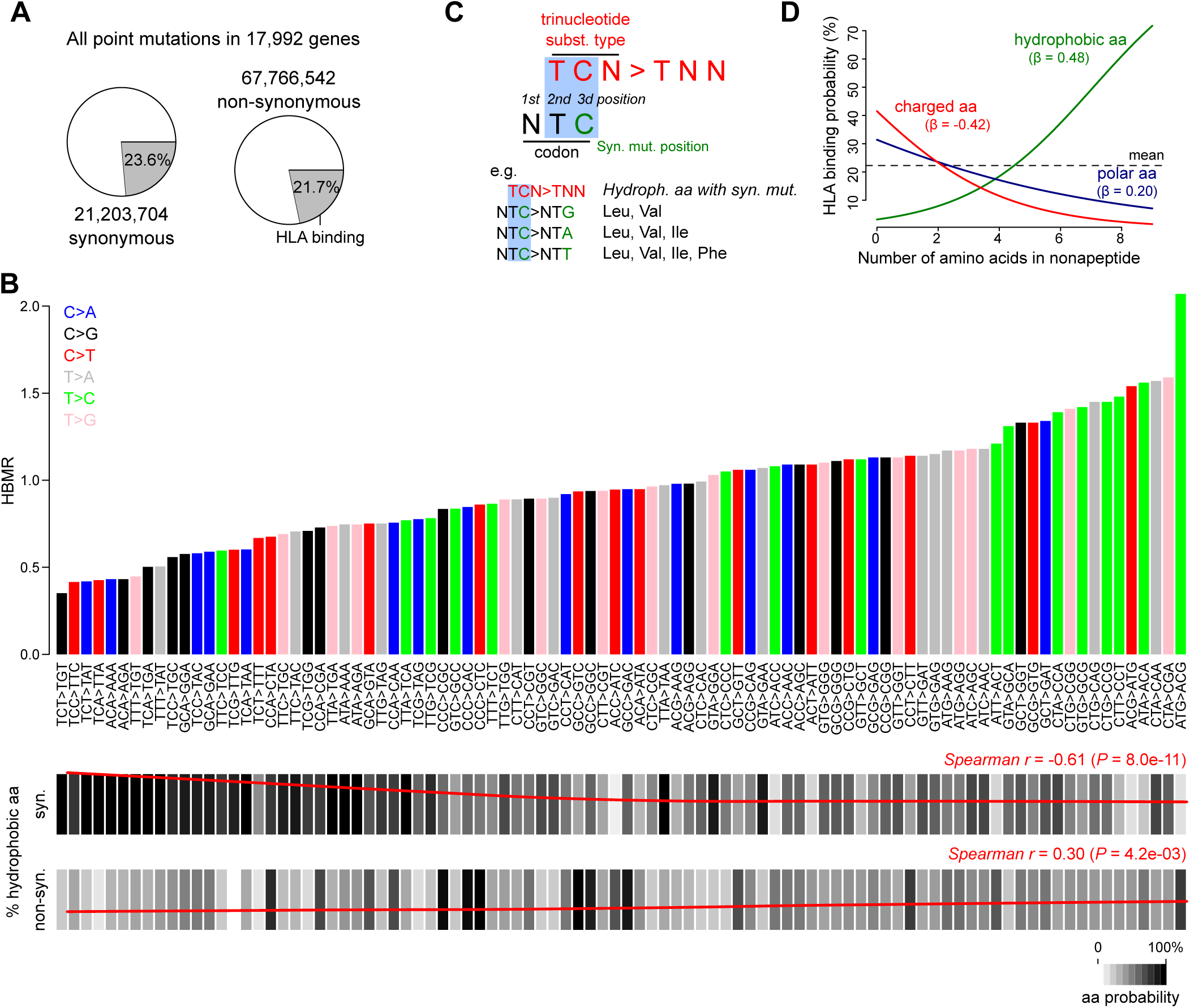
Influence of trinucleotide substitution types on HLA-binding properties. (**A**) All possible synonymous and non-synonymous mutations were determined in 17,992 genes. Pie charts indicate the proportions of these mutations that are located in HLA-binding regions. (**B**) Bar plot on top indicates HBMR values for each trinucleotide substitution type. Main substitution types are coloured as indicated by the legend on top left. Note that HBMR values are not derivable for 4 trinucleotide substitution types (ATT>AAT, ATT>AGT, ACT>AGT and ACT>AAT) due to the absence of synonymous mutations resulting from these substitution types (e.g. an ATT>AAT substitution can never be synonymous). Below the bar plot, the frequency of synonymous and non-synonymous mutations hitting hydrophobic amino acids is indicated for each substitution type (scale indicated on bottom right). Loess regression line in red with spearman correlation coefficient indicated on top right. (**C**) Illustration of TCN>TNN mutations mainly resulting in synonymous mutations in hydrophobic amino acid codons. (**D**) Logistic regression line indicating the correlation between a nonapeptide’s number of hydrophobic/charged/polar amino acids (0 to 9) and the HLA-binding probability. Regression coefficients (β) are given for each amino acid class.

### High synonymous mutation probabilities in hydrophobic amino acid codons correlate to lower perceived mutation rates in HLA-binding regions

We next aimed to explain this unexpected association between trinucleotide substitution types and HLA-binding properties. Because different sequence contexts imply different amino acid codon probabilities on the one hand, while different physicochemical properties of amino acids influence binding to HLAs on the other hand, we examined whether an effect through one of the 3 main amino acid classes (hydrophobic, polar or charged) could explain this association. Therefore, we determined whether there was a relationship between 1) the trinucleotide substitution types’ HBMR values and the class of the amino acid codon containing this mutation and 2) the number of amino acids from a certain class in a nonapeptide and its HLA-binding capacity.

We first focused on the correlation between HBMR values and the amino acid classes and determined which proportion of each substitution type’s synonymous and non-synonymous mutations occurred in the three amino acid class codons. A strong negative correlation was observed between a trinucleotide substitution type’s HBMR value and the proportion of synonymous mutations in hydrophobic amino acid codons (Spearman r = −0.61, *P* = 8.0e-11; Fig. 3B), while an opposite, though weaker positive correlation was noted for non-synonymous mutations hitting hydrophobic amino acid codons (Spearman r = 0.30, *P* = 4.2e-03; Fig. 3B). To better understand this association, we extended the correlation analysis to all 20 individual amino acids and observed that the hydrophobic amino acid effect was mainly attributed to Leu, Val and Iso (Fig. S6). Strikingly, all these hydrophobic amino are encoded by codons with a thymine on the second codon position (Fig. S7). Combined with the observation that most of the corresponding trinucleotide substitution types are from the format TCN>TNN, this association is explained by the upstream T of the substitution type matching with the T at the second codon position and the substituted nucleotide matching with the third codon position (Fig. 3C). Indeed, while any codon with a T at the second position is hydrophobic, any mutation involving the third position of a Leu or Val codon will always result in a synonymous mutation. This is also the case for most mutations hitting the same position in Ile and for some mutations at the Phe codon as exemplified in Fig. 3C.

As hydrophobic amino acids are known to influence HLA-binding affinities (33), we hypothesized that the association between trinucleotide substitution types and probabilities of (synonymous) mutations hitting hydrophobic amino acid codons could also explain the higher observed effects on the HLA-binding properties of the translated peptides. Therefore, we examined the effect of changing a nonapeptide’s number of hydrophobic amino acids on its HLA-binding capacity by randomly sampling from 1 million CDS regions and determining the translated peptides’ HLA-binding affinity. By applying a binomial logistic regression model using the number of amino acids as observed variable and the HLA-binding capacity (binding versus non-binding) as response variable, we could indeed demonstrate a strong positive association between the number of hydrophobic amino acids in a peptide and the HLA-binding capacity (regression coefficient *β* = 0.48, Fig. 3D).

These results demonstrate that certain trinucleotide substitution types, like TCN>TNN, which occur frequently in metastatic melanoma, bladder cancer and cervical cancer, are likely to lead to synonymous mutations hitting Leu, Val and Ile. Because these amino acids are also more frequent in HLA-binding peptides, this leads to higher synonymous mutation rates in these peptides, implying lower non-synonymous mutation rates when synonymous mutations are used as a background reference.

### Absence of neoantigen depletion signals after correcting for the trinucleotide substitution effect

Our results show that the main factor determining differential mutation rates between HLA binding and non-binding peptides are differences in trinucleotide substitution probabilities. We next aimed to determine whether any remaining trace of neoantigen depletion would be detectable after correcting for these trinucleotide effects. Therefore, for each cancer, we normalized the observed HBMR value to its expected HBMR value. The latter was calculated based on the (trinucleotide) substitution probabilities derived from each cancer’s aggregated mutational signature on the one hand and the HLA-binding annotation developed in this study on the other hand (see *Materials and Methods*). Using the resulting normalized HBMR, all putative signals of neoantigen depletion disappeared in our dataset, except for a limited signal in lung adenocarcinoma (Fig. 4). Similarly, also the dN/dS values in HLA-binding regions did not suggest any signal of negative selection after correcting for the trinucleotide substitution probability effect (Fig. S2B). Notably, correcting using mutation probabilities derived from models that do not take into account the adjacent sequence context, the corrected signals were (falsely) suggestive for neoantigen depletion in e.g. melanoma and bladder cancer.

**Figure 4.**
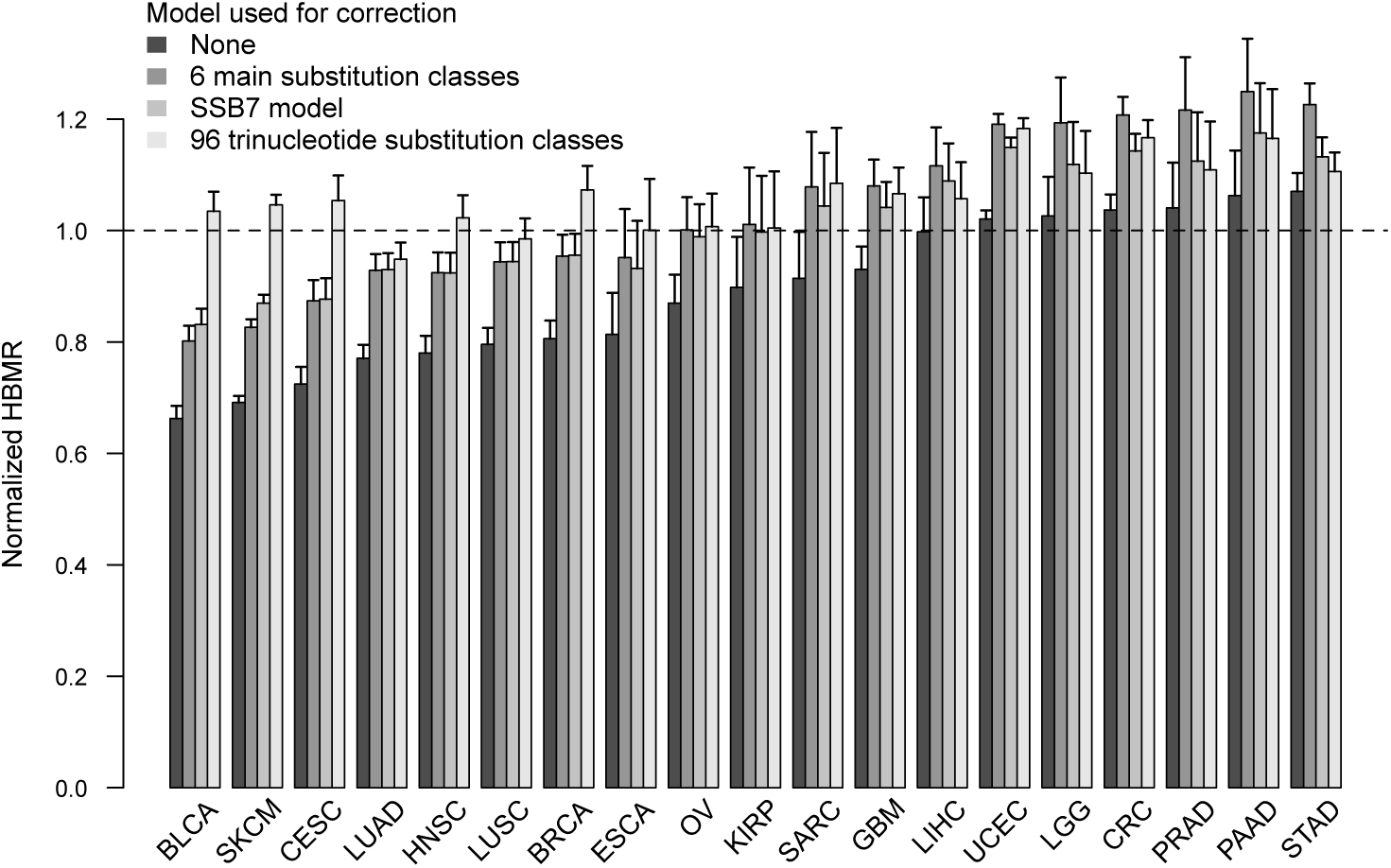
Absence of clear neoantigen depletion signals after correcting for trinucleotide-based mutational signature effects. Bar plot shows normalized HBMR values for 19 different cancer types. HBMR values were obtained by normalization of the observed HBMR values to the expected HBMR values. The latter values were calculated using mutation probabilities derived from different models as indicated on top left. Error bars indicate 95% confidence intervals, calculated using Fisher’s exact test. See table S1 for cancer type abbreviations.

These results confirm the absence of any detectable neoantigen depletion in cancer genomics data (with the possible exception of lung cancer) and emphasize the importance of using an accurate (trinucleotide-based) model to correct for mutational signature effects.

### Lower mutation rates in known epitopes are secondary to HLA-binding properties

Because our results suggest a lack of clear signals of neoantigen depletion based on the analysis of HLA-binding regions on the one hand, while lower-than-expected non-synonymous mutation rates and hence negative selection have been suggested in epitope sequences in cancer on the other hand (19), we next aimed to explain this discrepancy.

We mapped 66,536 human epitopes, derived from the immune epitope database (IEDB), to the human genome and compared non-synonymous mutation rates within and outside of mapped epitope regions (Fig. 5A). This approach has the additional advantage that it is not restricted to nonapeptides (Fig. 5A). To compare non-synonymous mutation rates between epitope and non-epitope mapped genomic regions, we defined an epitope mutation ratio (EMR), similar to the HBMR approach for HLA-binding regions. This EMR was significantly below 1 in metastatic melanoma (EMR = 0.89, *P* = 3.8e-10) and cervical cancer (EMR = 0.91, *P* = 0.018) (Fig. S8). However, similar results were obtained using the simulation approach (EMR = 0.89 and 0.96 for melanoma and cervical cancer respectively; Fig. S8), again suggesting that selection was not causing these lower mutation rates.

**Figure 5.**
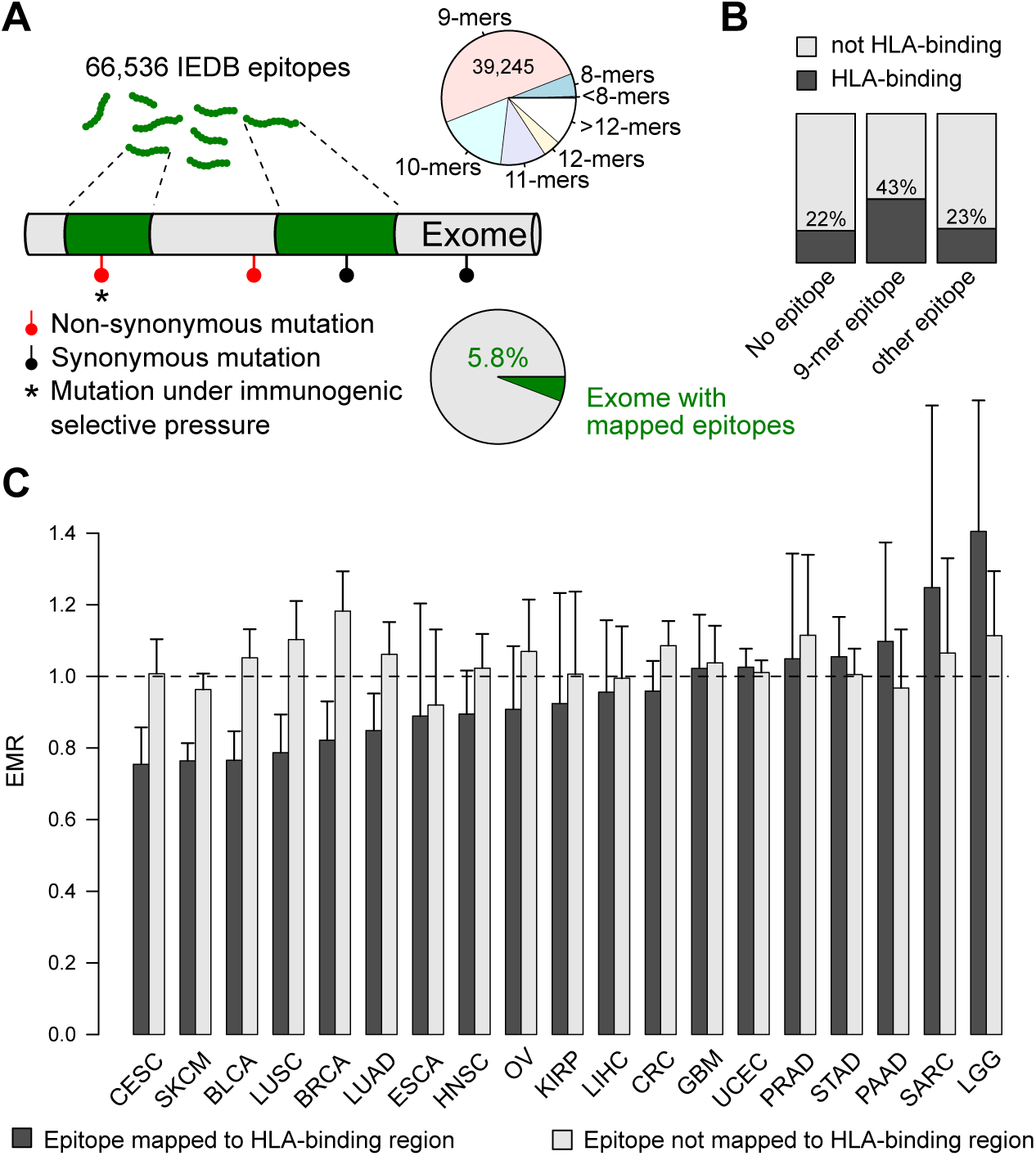
Analysis of negative selection signals in IEDB epitopes. (**A**) 66,536 IEDB epitopes were mapped to the human genome and mutation rates were compared between mutations within and outside of the epitope regions. Pie chart on top and bottom indicate the sizes of the mapped epitopes and the proportion of the exome containing mapped epitopes respectively. (**B**) Overlap between epitope regions and HLA-binding regions. Only 9-mers showed a clear overlap with the HLA-binding regions. (**C**) Bar plot compares the epitope mutation ratios (EMR, i.e. ratio of n/s in epitope to non-epitope regions) between HLA-binding and non-binding regions for different cancers. Error bars indicate 95% confidence intervals, calculated using Fisher’s exact test.

We observed a considerable overlap between 9-mer epitope-mapped genomic regions and the HLA-binding regions described higher (Fig. 5B). To determine whether the lower (observed and simulated) mutation rates in melanoma and cervical cancer were caused by this overlap, we repeated the analysis while separating epitopes mapped to HLA-binding regions from epitopes mapped to non-binding regions. This analysis showed that lower mutation rates in epitope-mapped regions were indeed completely attributed to the HLA-binding properties of the epitopes (Fig. 5C). Mutation rates in epitopes mapped to HLA-binding were very similar to the overall results using HLA-binding properties as shown in Fig. 2C.

## DISCUSSION

In this study we observed an apparent reduction of somatic mutations occurring in genomic regions encoding HLA-binding nonapeptides in several cancers. While this effect has been attributed to negative selection of immunogenic mutations (10, 19), we demonstrated that these lower mutation rates were mainly determined by the correlation between different mutational signatures (i.e. different trinucleotide substitution type probabilities) and the resulting peptides’ HLA affinities, rather than neoantigen depletion, and showed that the main mediator is hydrophobicity of the amino acids forming the peptide.

We focussed on somatic mutations located in genomic regions that can be translated to HLA-binding peptides. As these peptides can be presented to CTLs, the assumption was that some of these mutations change CTL contact residues, leading to neoantigen formation and hence CTLl reactivity. The resulting CTL-mediated cancer cell killing is expected to lead to negative selection of these mutations and hence neoantigen depletion. To detect signals of this neoantigen depletion, we first annotated the complete human genome for its HLA-binding capacity, where HLA-binding regions are defined as those regions in the genome that are translatable to HLA-binding nonapeptides. We then compared the non-synonymous mutation rates in HLA-binding to non-binding peptides. While our results apparently suggested strong signals of neoantigen depletion in several cancers known to be responsive to immunotherapy (7, 8, 34, 35), strikingly and surprisingly, almost completely identical results were obtained when a simulated mutation dataset was used, upon which (negative) selection could not have acted. We demonstrated that this effect could be mainly attributed to different substitution types correlating to the HLA-binding properties of the translated peptides. For the substitution types with the lowest mutation rates in HLA-binding regions, a remarkably high frequency of TCN>TNN mutations was observed. As this implies a thymine on the second codon position in case of synonymous mutations, synonymous mutations will mainly hit hydrophobic amino acid codons. This will result in low apparent non-synonymous mutation rates when synonymous mutations are used as a background reference. As we also showed that the number of hydrophobic amino acids are a major determinant of a peptide’s HLA-binding capacity, cancers with more TCN>TNN or related substitutions (e.g. bladder cancer and melanoma) will contain more synonymous mutations in hydrophobic amino acid codons and hence also HLA-binding regions, which is ultimately perceived as lower non-synonymous mutation rates, explaining our findings.

Recent studies have shown that standard selection metrics such as dN/dS can be confounded when not taking mutational signatures into account (22, 23). The key finding of this study is that also HLA-binding affinities are influenced by different mutational signatures, and should be corrected for. In this context we also showed that any apparent signal of negative selection in known IEDB epitopes is secondary to the HLA-binding affinities of the epitopes and hence related to different mutational signatures rather than negative selection. Our results contradict the results of from a recent report describing negative selection in HLA-binding epitope regions (19). The main reason for this contradiction seems to be the intrinsic lack of correcting for TCN>TNN mutation probabilities by the model that was used in that study.

Neoantigen depletion is a generally accepted immune evasion mechanism for the evolving cancer cell. Our results indicate that signals of neoantigen depletion are overall weak to absent in the untreated cancer genome. While we cannot exclude that this is related to poor accuracy to predict HLA affinities and neoantigen formation, it is remarkable that signals of negative selection in general are weak in cancer mutation data (22, 23, 36, 37). As neoantigen depletion results from a specific form of negative selection, it should in the end also be detectable as a reduction in overall mutation rates in cancers that are known to be influenced by the immune system, even without considering HLA affinities. As this is not the case, the actual absence of neoantigen depletion, rather than technical reasons, could explain our results. Therefore, we hypothesize that more efficient immune evasion mechanism (e.g. *HLA* loss, *PDL1* amplification, …) than neoantigen depletion are active early during tumour evolution, before negative selection manages to shape the cancer genome. If this is indeed the case, signals of neoantigen depletion are only expected in the absence of these escape mechanisms, such as after ICB therapy. Interestingly, ICB therapy has indeed been suggested to result in neoantigen depletion post-therapy (20).

## Supporting information

## ACKNOWLEDGEMENTS

The results published here are in whole or part based upon data generated by TCGA. Information about TCGA and the investigators and institutions who constitute the TCGA research network can be found at http://cancergenome.nih.gov. We are most grateful to the patients, investigators, clinicians, technical personnel, and funding bodies who contributed to TCGA, thereby making this study possible.

## SUPPLEMENTARY TABLE AND FIGURES LEGENDS

**Figure S1. HLA affinity analysis of the most frequent HLA alleles** (**A**) Comparison of HLA allele frequencies in TCGA and the general population. HLA alleles for TCGA samples were calculated using Polysolver (11). Population frequencies were derived from a US Caucasian blood donor population, downloaded from Allele frequency net (28). The 2 most frequent alleles are indicated for each HLA gene. Pearson correlation coefficients and *P* values indicated on top left. (**B**) Spearman correlation plot between the predicted nonapeptide Kd values of the most frequent HLA alleles and the aggregated Kd as indicated. Correlations were determined based on 1 million random coding regions. (**C**) Comparison of the proportion of the HLA-binding exome for each individual allele (as indicated) and after aggregating the affinities of the 6 alleles (shown on top right).

**Figure S2. dN/dS in HLA-binding and non-binding peptides for different cancers** Bar plots show dN/dS values for HLA-binding and non-binding peptides as indicated. Error bars indicate 95% confidence intervals. The ratio dN/dS was calculated considering different trinucleotide substitution probabilities (see *Materials and Methods*) without (**A**) or with correction (**B**) for HLA-binding regions. See table S1 for cancer type abbreviations.

**Figure S3. HBMR calculations using alternative approaches to determine HLA-binding regions** Similar analysis as shown in main Fig. 2A-C, but using alternative approaches to determine HLA-binding regions: 1) Sample-specific alleles (*top panels*): Aggregated HLA-affinities were calculated using the TCGA sample-specific HLA alleles as determined by Polysolver. 2) Pooled HLA alleles (*middle panels*): HLA affinities determined for the 6 different HLA alleles were pooled together and considered as separate mutations. 3) Kd cut-off 50nM (*bottom panels*): a more stringent Kd cut-off of 50nM was used to consider nonapeptides HLA-binding. Note that while the proportions of mutations hitting HLA-binding regions was lower using the pooled HLA or more stringent Kd cut-off approach, the observed effect on HBMR did not change substantially.

**Figure S4. Absence of selection signals in the simulated mutation database** The ratio of the number of non-synonymous mutations over the number of synonymous mutations was calculated for each gene from the observed and simulated mutation database. Dashed line indicates the median n/s ratio from the observed mutations. Genes are ranked based on the observed n/s ratio and only the top 50 ranked genes are shown.

**Figure S5. Correlation between HBMR values and the aggregated mutational signatures of 19 analysed cancer types** (**A**) Bar plots show the proportion of 96 different trinucleotide substitution types for each cancer as indicated. Substitution types are ranked based on increasing HBMR values. See Fig. 3B for HBMR details and colour legends. (**B**) Correlation between the proportion of TCN>TNN mutations and HBMR values for each analysed cancer as indicated. Pearson correlation coefficient and *P* value indicated on top left. See table S1 for cancer type abbreviations.

**Figure S6. Correlation between HBMR values of different trinucleotide substitution types and their targeted amino acid codons** The probability of synonymous (*top panel*) and non-synonymous (*bottom panel*) mutations hitting individual amino acids and amino acid classes is indicated for each substitution type (scale indicated on bottom right). Significant Spearman correlations (*P* below 0.05) are shown for each amino acid (scale indicated on bottom left). Loess regression line for amino acid classes in red. Trinucleotide substitution types are ordered on increasing HBMR values. See main Fig. 3B for HBMR details.

**Figure S7. Genetic code** Overview of the genetic code. Amino acid classes (hydrophobic/polar/charged) are coloured as indicated.

**Figure S8. Analysis of mutation rates in IEDB epitopes** 66,536 IEDB epitopes were mapped to the human genome and mutation rates were compared between mutations within and outside of the epitope regions. See Fig. 5A for details. Bar plot compares EMRs (Epitope mutation rate, i.e. the ratio of n/s in epitope to non-epitope regions) between different cancers and calculated using the observed and simulated mutation data as indicated. Error bars indicate 95% confidence intervals, calculated using Fisher’s exact test.

**Table S1 Overview of analysed TCGA mutation data**

