## Supplementary material for "Lack of detectable neoantigen depletion in the untreated cancer genome"

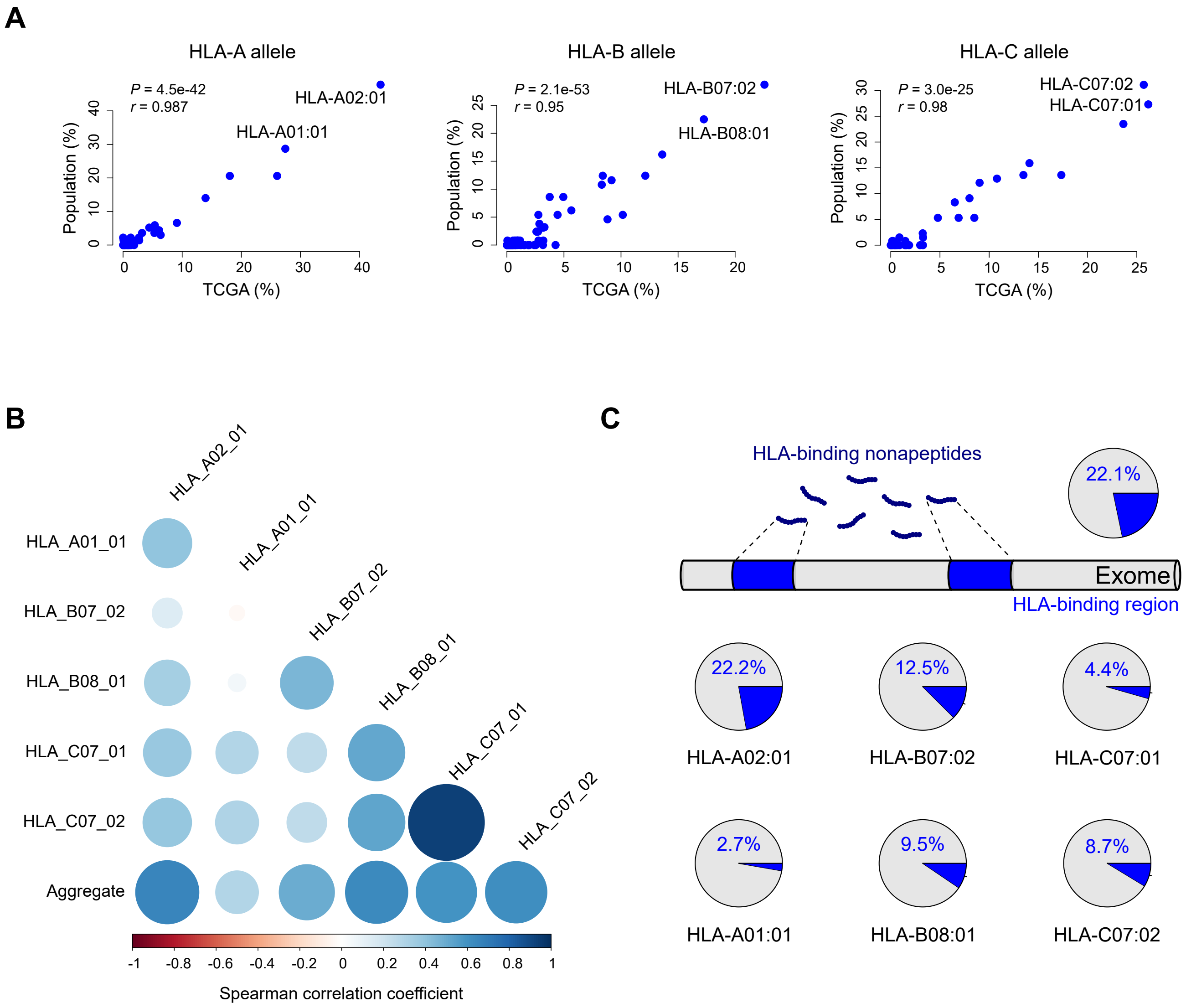
A
HLA-A allele
P = 4.5e-42
r = 0.987
HLA-A02:01
HLA-A01:01
Population (%)
TCGA (%)
HLA-B allele
P = 2.1e-53
r = 0.95
HLA-B07:02
HLA-B08:01
Population (%)
TCGA (%)
HLA-C allele
P = 3.0e-25
r = 0.98
HLA-C07:02
HLA-C07:01
Population (%)
TCGA (%)
B
HLA_A02_01
HLA_A01_01
HLA_B07_02
HLA_B08_01
HLA_C07_01
HLA_C07_02
Aggregate
-1 -0.8 -0.6 -0.4 -0.2 0 0.2 0.4 0.6 0.8 1
Spearman correlation coefficient
C
HLA-binding nonapeptides
22.1%
Exome
HLA-binding region
22.2%
12.5%
4.4%
HLA-A02:01
HLA-B07:02
HLA-C07:01
2.7%
9.5%
8.7%
HLA-A01:01
HLA-B08:01
HLA-C07:02

*Suppl. fig. 1*

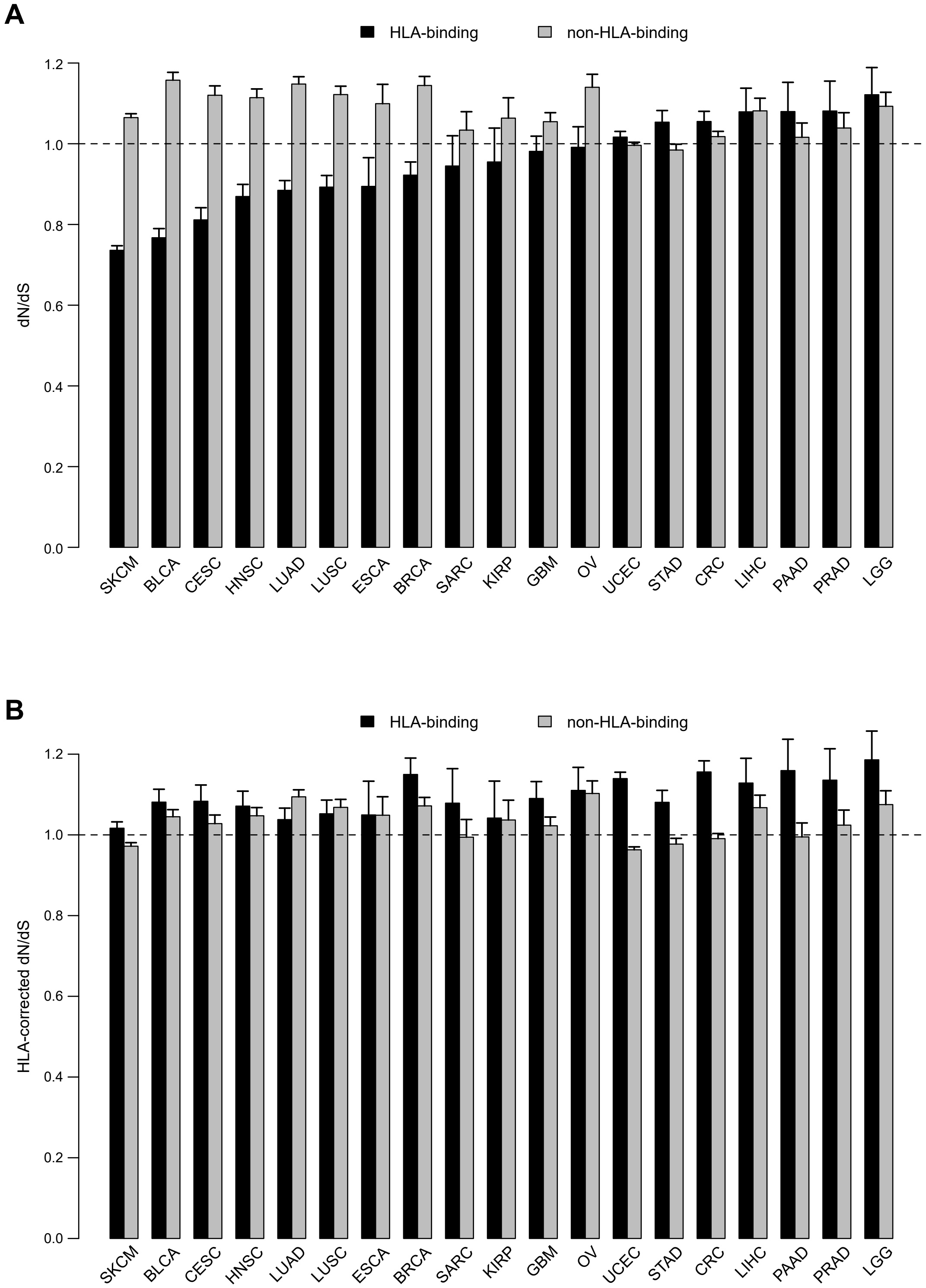
A
HLA-binding
non-HLA-binding
dN/dS
1.2
1.0
0.8
0.6
0.4
0.2
0.0
SKCM
BLCA
CESC
HNSC
LUAD
LUSC
ESCA
BRCA
SARC
KIRP
GBM
OV
UCEC
STAD
CRC
LIHC
PAAD
PRAD
LGG
B
HLA-binding
non-HLA-binding
HLA-corrected dN/dS
1.2
1.0
0.8
0.6
0.4
0.2
0.0
SKCM
BLCA
CESC
HNSC
LUAD
LUSC
ESCA
BRCA
SARC
KIRP
GBM
OV
UCEC
STAD
CRC
LIHC
PAAD
PRAD
LGG

*Suppl. fig. 2*

### Sample-specific HLA alleles

| | HLA binding - | HLA binding + | % |
|---|---|---|---|
| s | 408,170 | 117,507 | 22.4% |
| n | 1,043,513 | 267,179 | 20.4% |
| n/s | 2.56 | 2.27 | |
| HBMR | 0.89 | | |

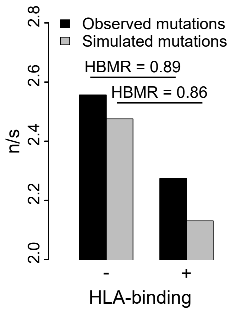


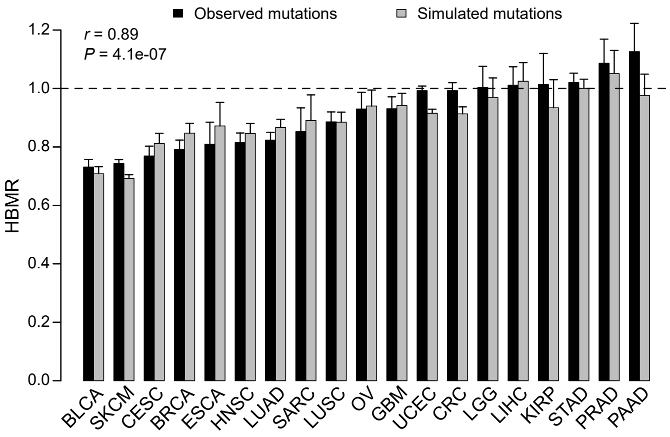


### Pooled HLA alleles

| | HLA binding - | HLA binding + | % |
|---|---|---|---|
| s | 2,814,964 | 339,098 | 10.7% |
| n | 7,062,565 | 801,587 | 10.2% |
| n/s | 2.51 | 2.36 | |
| HBMR | 0.94 | | |

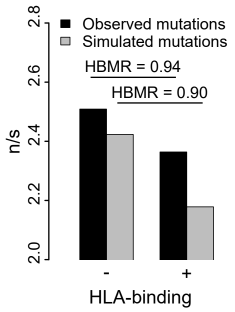


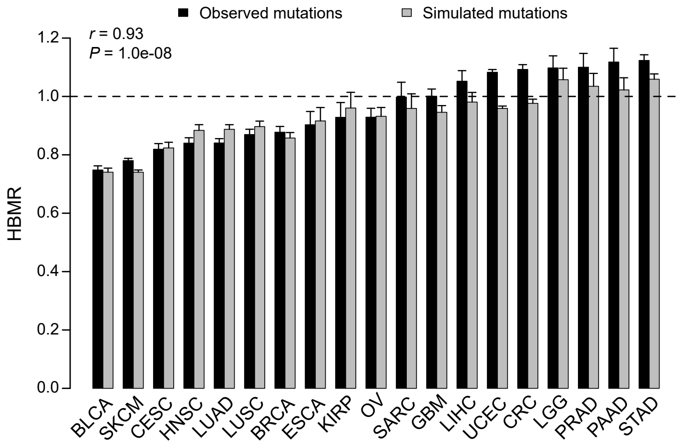


### Kd cut-off 50nM

| | HLA binding - | HLA binding + | % |
|---|---|---|---|
| s | 507,734 | 17,943 | 3.4% |
| n | 1,273,291 | 37,401 | 2.9% |
| n/s | 2.51 | 2.08 | |
| HBMR | 0.83 | | |

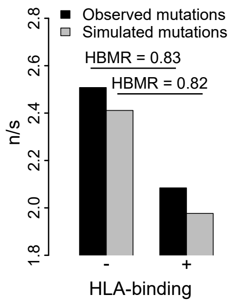


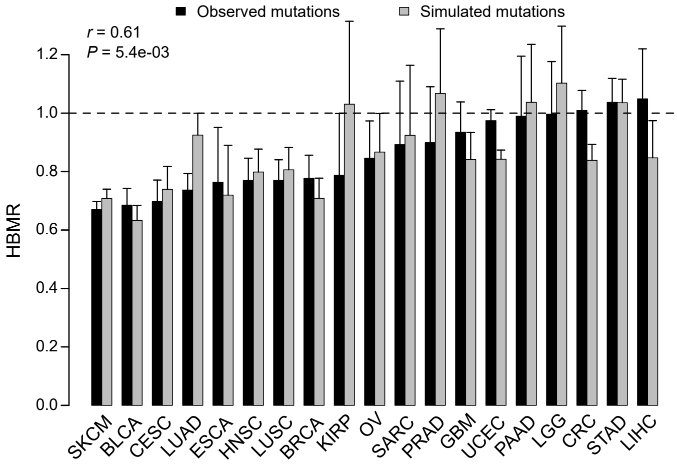


*Suppl. fig. 3*

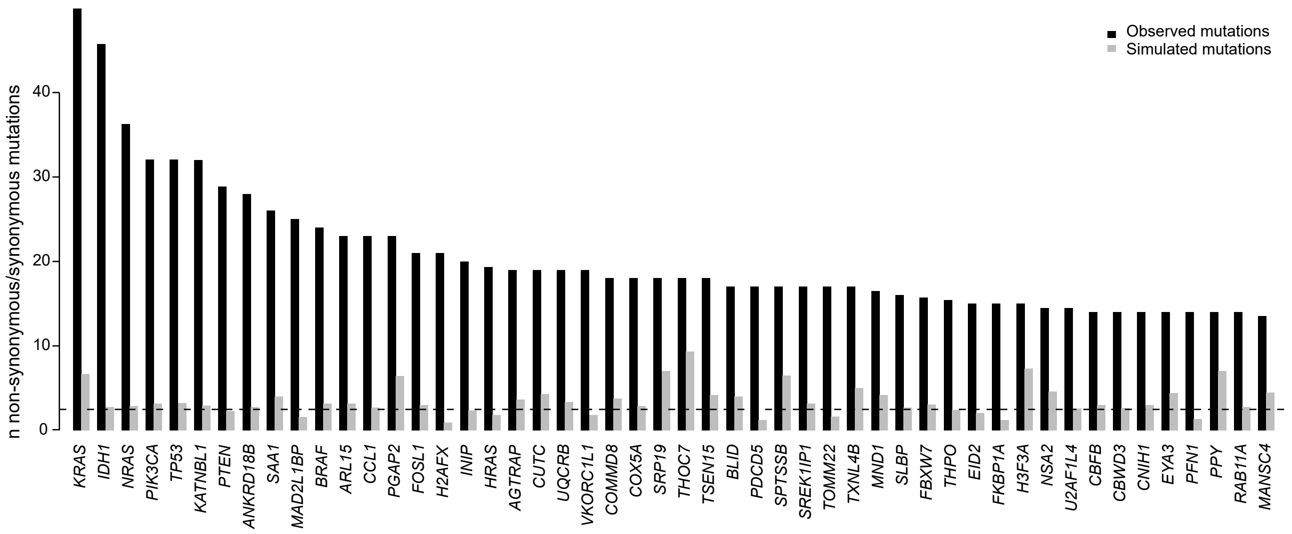
Observed mutations
Simulated mutations
n non-synonymous/synonymous mutations
0
10
20
30
40
KRAS
IDH1
NRAS
PIK3CA
TP53
KATNBL1
PTEN
ANKRD18B
SAA1
MAD2L1BP
BRAF
ARL15
CCL1
PGAP2
FOSL1
H2AFX
INIP
HRAS
AGTRAP
CUTC
UQCRB
VKORC1L1
COMMD8
COX5A
SRP19
THOC7
TSEN15
BLID
PDCD5
SPTSSB
SREK1IP1
TOMM22
TXNL4B
MND1
SLBP
FBXW7
THPO
EID2
FKBP1A
H3F3A
NSA2
U2AF1L4
CBFB
CBWD3
CNIH1
EYA3
PFN1
PPY
RAB11A
MANSC4

*Suppl. fig. 4*

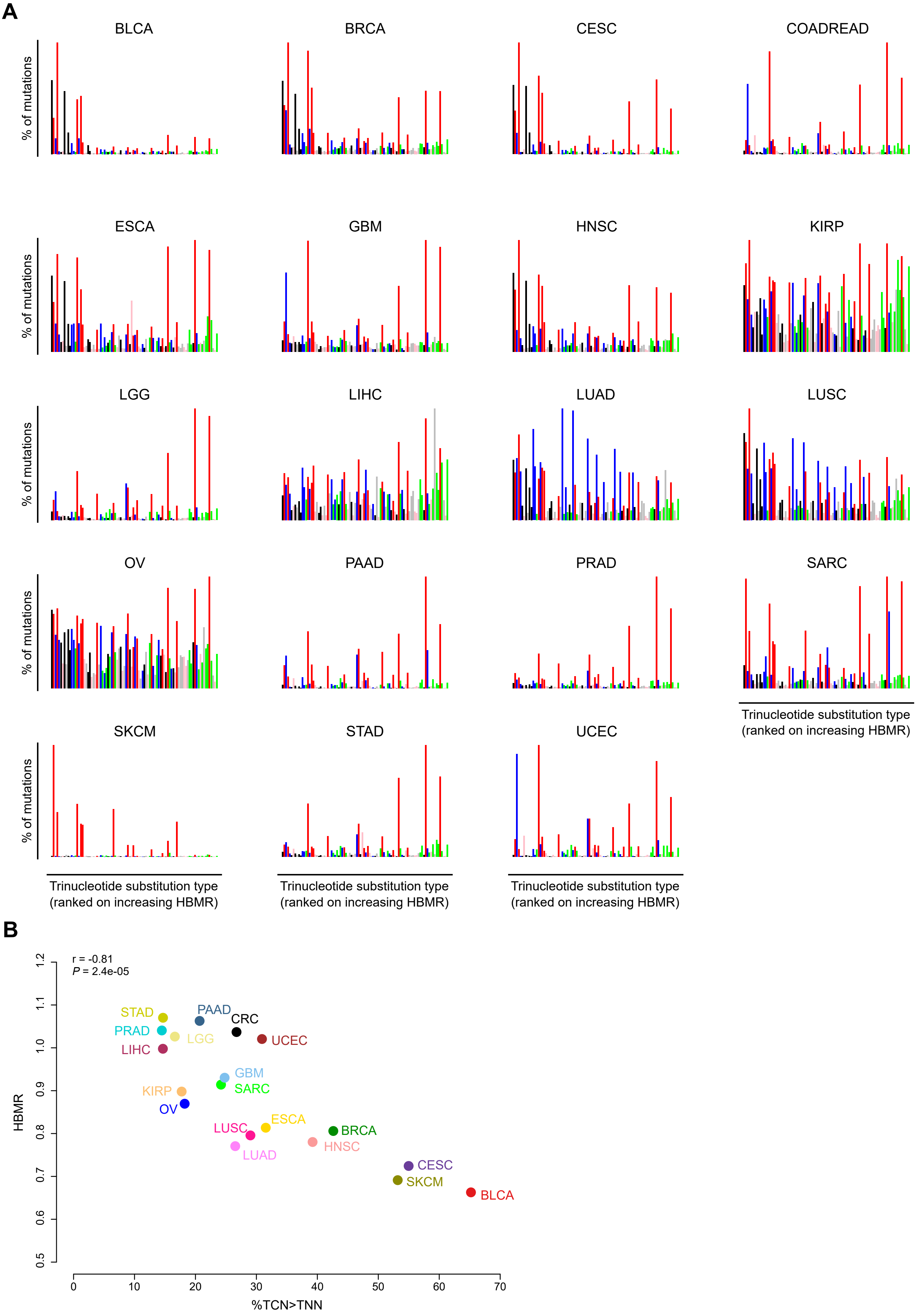

A
BLCA
BRCA
CESC
COADREAD
ESCA
GBM
HNSC
KIRP
LGG
LIHC
LUAD
LUSC
OV
PAAD
PRAD
SARC
SKCM
STAD
UCEC
% of mutations
Trinucleotide substitution type (ranked on increasing HBMR)
B
r = -0.81
P = 2.4e-05
HBMR
%TCN>TNN
0.5
0.6
0.7
0.8
0.9
1.0
1.1
1.2
0
10
20
30
40
50
60
70
STAD
PAAD
CRC
PRAD
LGG
UCEC
LIHC
GBM
SARC
KIRP
OV
LUSC
ESCA
BRCA
HNSC
LUAD
CESC
SKCM
BLCA


*Suppl. fig. 5*

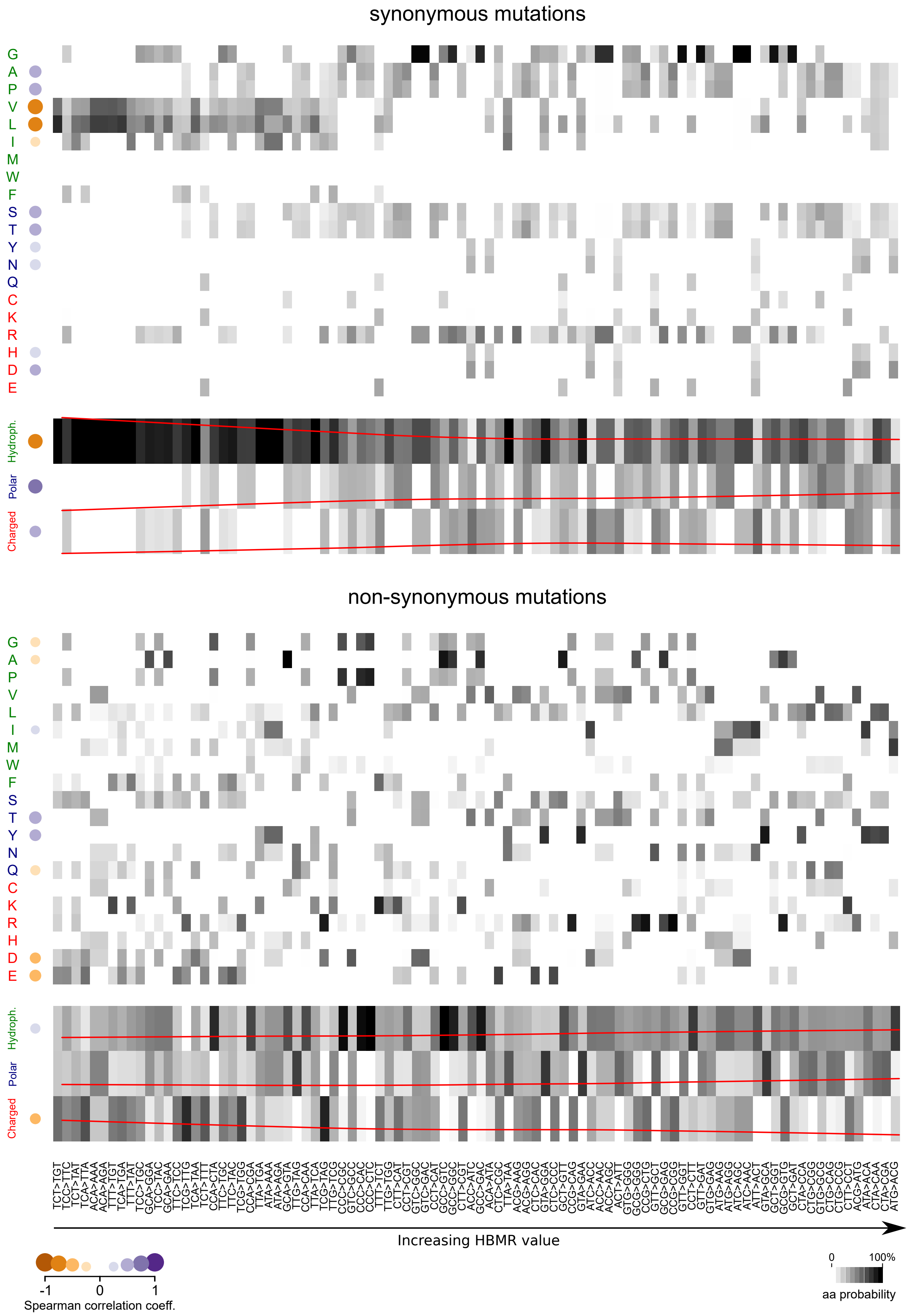
synonymous mutations
G
A
P
V
L
I
M
W
F
S
T
Y
N
Q
C
K
R
H
D
E
Hydroph.
Polar
Charged
non-synonymous mutations
Increasing HBMR value
-1
0
1
Spearman correlation coeff.
0
100%
aa probability

*Suppl. fig. 6*

Second column

| First column | T | C | A | G | Third column |
|---|---|---|---|---|---|
| T | *TTT* F | *TCT* S | *TAT* Y | *TGT* C | T |
| | *TTC* F | *TCC* S | *TAC* Y | *TGC* C | C |
| | *TTA* L | *TCA* S | *TAA* stop | *TGA* stop | A |
| | *TTG* L | *TCG* S | *TAG* stop | *TGG* W | G |
| C | *CTT* L | *CCT* P | *CAT* H | *CGT* R | T |
| | *CTC* L | *CCC* P | *CAC* H | *CGC* R | C |
| | *CTA* L | *CCA* P | *CAA* Q | *CGA* R | A |
| | *CTG* L | *CCG* P | *CAG* Q | *CGG* R | G |
| A | *ATT* I | *ACT* T | *AAT* N | *AGT* S | T |
| | *ATC* I | *ACC* T | *AAC* N | *AGC* S | C |
| | *ATA* I | *ACA* T | *AAA* K | *AGA* R | A |
| | *ATG* M | *ACG* T | *AAG* K | *AGG* R | G |
| G | *GTT* V | *GCT* A | *GAT* D | *GGT* G | T |
| | *GTC* V | *GCC* A | *GAC* D | *GGC* G | C |
| | *GTA* V | *GCA* A | *GAA* E | *GGA* G | A |
| | *GTG* V | *GCG* A | *GAG* E | *GGG* G | G |

Hydrophobic amino acid (codon)
Polar amino acid (codon)
Charged amino acid (codon)

*Suppl. fig. 7*

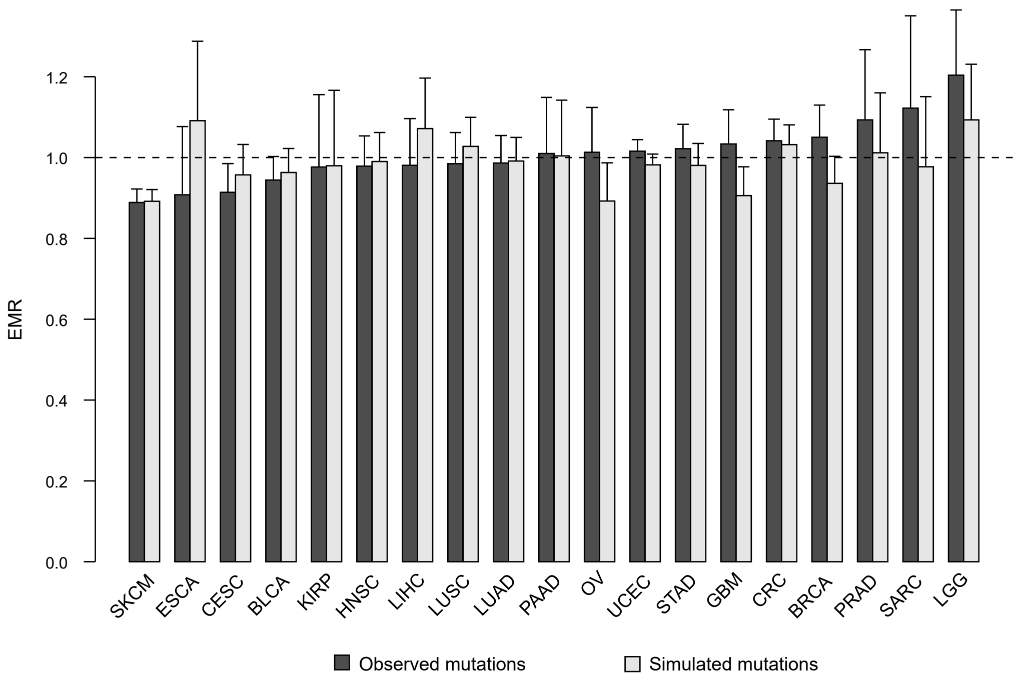

1.2
1.0
0.8
0.6
0.4
0.2
0.0
EMR
SKCM
ESCA
CESC
BLCA
KIRP
HNSC
LIHC
LUSC
LUAD
PAAD
OV
UCEC
STAD
GBM
CRC
BRCA
PRAD
SARC
LGG
Observed mutations
Simulated mutations


*Suppl. fig. 8*

| TCGA abbreviation | cancer type | n samples | n mutations |
|---|---|---|---|
| ACC | Adrenocortical carcinoma | 87 | 4232 |
| BLCA | Bladder Urothelial Carcinoma | 390 | 93464 |
| BRCA | Breast invasive carcinoma | 904 | 70633 |
| CESC | Cervical squamous cell carcinoma and endocervical adenocarcinoma | 281 | 60202 |
| CHOL | Cholangiocarcinoma | 33 | 1794 |
| CRC | Colorectal cancer | 426 | 161449 |
| DLBC | Lymphoid Neoplasm Diffuse Large B-cell Lymphoma | 37 | 4318 |
| ESCA | Esophageal carcinoma | 124 | 14333 |
| GBM | Glioblastoma multiforme | 388 | 61331 |
| HNSC | Head and Neck squamous cell carcinoma | 488 | 70172 |
| KICH | Kidney Chromophobe | 9 | 164 |
| KIRC | Kidney renal clear cell carcinoma | 84 | 3793 |
| KIRP | Kidney renal papillary cell carcinoma | 222 | 12196 |
| LAML | Acute Myeloid Leukemia | 138 | 7028 |
| LGG | Brain Lower Grade Glioma | 502 | 26100 |
| LIHC | Liver hepatocellular carcinoma | 312 | 31768 |
| LUAD | Lung adenocarcinoma | 407 | 112204 |
| LUSC | Lung squamous cell carcinoma | 306 | 80066 |
| MESO | Mesothelioma | 80 | 2482 |
| OV | Ovarian serous cystadenocarcinoma | 333 | 36369 |
| PAAD | Pancreatic adenocarcinoma | 145 | 21331 |
| PCPG | Pheochromocytoma and Paraganglioma | 176 | 1745 |
| PRAD | Prostate adenocarcinoma | 434 | 19217 |
| SARC | Sarcoma | 216 | 13647 |
| SKCM | Skin Cutaneous Melanoma | 466 | 302761 |
| STAD | Stomach adenocarcinoma | 390 | 118218 |
| TGCT | Testicular Germ Cell Tumors | 144 | 2093 |
| THCA | Thyroid carcinoma | 414 | 4477 |
| THYM | Thymoma | 110 | 2221 |
| UCEC | Uterine Corpus Endometrial Carcinoma | 506 | 488042 |
| UCS | Uterine Carcinosarcoma | 51 | 7105 |
| UVM | Uveal Melanoma | 80 | 1414 |
